# New Quantitative Cost-Impact Effectiveness Indexes to Assist in Publication Decisions by Researchers in the Open Access Era

**DOI:** 10.1101/2025.10.10.681750

**Authors:** Julia C. Hardy, Christine Kim Vu, Dong Wang

## Abstract

Scientific publications have become the backbone of scientific communication since their foundation in 1665. The three main models for publishing are Traditional (or subscription-based), Open Access (OA), and Hybrid. As of July 1, 2025, the NIH requires that Author Accepted Manuscripts resulting from NIH-funded research be immediately publicly available. To comply with this new requirement, authors may be forced to pay an Article Processing Charge (APC) to publish Open Access, ranging from ∼$2000 to ∼$13,000 per article. With this change to the scientific publishing landscape, publishing costs shift from subscribers to authors causing authors to re-evaluate how they choose which journal to publish in. Here we analyze 75 popular biomedical journals to evaluate the publishing costs compared to the scientific impact (i.e. Impact Factor, CiteScore, SNIP) illustrated by three different Cost-Impact Effectiveness (CIE) metrics (APC/IF, APC/CS and APC/SNIP). To complement the new open access policy, our goal is to provide a resource to help the scientific community evaluate the impact-based cost effectiveness of different Open Access options during their journal selection process.

## Introduction

Scientific research is the driving force for numerous biomedical and technological advances. As the major communication platform for scientists, scientific publications are crucial for sharing cutting-edge discoveries and knowledge to the scientific community at large. The world’s first and longest-running scientific journal, *Philosophical Transactions*, was first launched in 1665 (1). Over the past 360 years, the academic publishing process has continuously evolved.

Nowadays, there are three main publishing models of journals: Traditional (or subscription-based), Open Access (OA), and Hybrid (2, 3). In the Traditional, subscription-based model, articles can only be accessed by subscribers of the journal or by users who pay per article. Alternatively, the Open Access model allows free public access to research articles. The hybrid model offers both subscription-based and open access publishing options through the journal.

There are different ‘routes’ to open access publishing. Under “Gold Open Access,” the final published version is made freely available immediately on the journal’s website but typically requires authors to pay an Article Processing Charge (APC). Many hybrid journals allow authors to choose if they would like to publish through subscription-based or “Gold Open Access” publishing avenues. Some journals also allow “Green Open Access,” also known as self-archiving, that allows authors to deposit an accepted version (submitted manuscript, author accepted manuscript (AAM), or the final published pdf) into a repository, often with an embargo time. Most subscription-based publishing release accepted manuscripts to the public after a 6 or 12-month embargo period, allowing authors to publicly share their accepted manuscripts without paying an APC (4, 5).

Over the past two decades, open access publishing has grown rapidly, with many journals shifting from subscription-based to open access models (4, 6). While readers benefit from free access, the cost burden has shifted to authors through often substantial APCs (7). In promoting transparency in research, funding agencies and institutions-including HHMI (8), Wellcome Research (9), Max-Planck-Gesellschaft Society (10), University of California (11), and NIH (12) – have Public Access Policy mandates that Author Accepted Manuscripts to be freely available for public access. Notably, NIH requires that Author Accepted Manuscripts resulting from NIH-funded research be immediately publicly available as of July 1, 2025 (13, 14).

Researchers now face mounting financial stress from two fronts: the uncertainty of budget cutting of grants and an increase of publication costs. Indeed, open access publication costs could become an unbearable burden and a much more significant portion of total research. In the last year, open access only journals have raised their APC by 6.5% with a maximum APC of $8,900 and hybrid journals have raised their APC by 3% with a maximum APC of $12,690 (15). Several major publishers are indicating that if an author has a zero-embargo mandate, the authors will be forced to select “Gold Open Access” publishing and therefore pay the APC. Other publishers are considering charging a fee to deposit the AAM without an embargo. The shift to open access publishing will inevitably change how authors evaluate journals for their publication choice. In addition to considerations on where to publish (e.g. disciplinary preference, scientific reputation of journals, time for peer-review processes), researchers will also need to take the publication cost as another critical factor into consideration during scientific publication (16, 17).

NIH recognizes that peer-reviewed publishing routes may result in publication costs, including but not limited to APCs, and provides some general “Points to Consider” for researchers and institutions in assessing reasonable costs for publication in relation to NIH award (18). In addition, NIH guidance clarifies that submitting Author Accepted Manuscripts to PubMed Central is free and that journals or publishers should not require a fee solely to deposit those manuscripts (19). Certain types of publisher charges tied to NIH-mandated public access are explicitly identified as unallowable under NIH cost principles (18, 20).

The NIH performed two analyses to evaluate recent publication costs using publicly available data from the Directory of Open Access Journals (7) and the United States Treasury Department. The first analysis looked at 7,350 journals worldwide and found the average APC is $1235.51 ($0.01 to $8,900) with a median of $950 (20). Looking at the 598 journals published in the United State, the average APC was $2,176.84 and a median of $2,040 (20). For the second analysis, the NIH looked at more than 1500 R01 grants awarded in Fiscal Year (FY) 2025 as of July 8, 2025 to evaluate the publication costs requested by NIH applicants (20). This showed that publication costs ranged from $0 to $12,000 and estimated that the average cost requested per publication was $2,565.07 to $3,104.06 (20). Moreover, NIH leadership has signaled further action to limit the financial burden on researchers and taxpayers by proposing a cap on allowable publication costs beginning in FY2026, and NIH has solicited input on how to maximize research funds by limiting allowable publishing costs (21). Unfortunately, there is a lack of quantitative metrics to evaluate the cost and impact of each publication, which would assist researchers with publication choice.

Here we propose three Cost-Impact Effectiveness (CIE) metrics that take into consideration both the APC and the journal’s impact, APC/IF, APC/CS and APC/SNIP, respectively. We hope that these new CIE metrics can assist researchers in choosing target journals in a cost-impact effective way.

### Impact-Based Cost Effectiveness Metrics of Journals

A journal’s scientific impact is usually evaluated by looking at how often it is referenced in other scientific works. The most common journal impact metric is the Journal Impact Factor by Clarivate based on data from Web of Science. The Journal Impact Factor (IF) is defined as all citations to a journal in the current year to publications from the previous two years compared to scholarly items (articles, reviews, and proceedings papers) published in the previous two years (22). Therefore, a higher IF usually indicates that the journal has a higher impact on the scientific community.

To better understand whether a journal’s scientific impact justifies its publishing cost, we first surveyed 75 representative biomedical journals and explored the relationship between APC and IF (Figure 1A, Supplemental Tables 1-3). Unfortunately, eLife did not receive an impact factor or CiteScore metric in 2025, so it is not included in the APC/IF and APC/CS metrics. Of the 75 journals, 70 (93.3%) have “Gold Open Access” options, with 48 (64%) have hybrid publishing options and 21 (28%) are open access only. The open access journals have a wide distribution of APCs ranging from ∼$2000 to ∼$13,000 per manuscript, except for the 6 (8%) subscription-only journals, such as *Science*, that have no APC. Additionally, 4 (5.3%) subscription-only journals offer embargo-free “Green Open Access.” Of our 75 surveyed journals, the median of APC for open access journals is $5790. To further assess the CIE metrics of each journal, we calculated the impact-normalized APC cost (APC/IF). APC/IF has an inverse relationship with impact-based cost effectiveness (Figure 1A and Supplemental Table 1). We then plotted the impact-normalized APC cost (APC/IF) vs IF (Figure 1B). The APC/IF ranges from $0 to $2174.42, with the median at $561 and $303 (for 25% percentile) among 75 journals we surveyed. To further analyze the pattern of APC/IF, we divided APC/IF into five bins based on IF values (Figure 1C). Interestingly, we observed the increasing IF-normalized APC medians and ranges with the decreasing IF values (Figure 1C). For the bin 1 journals (IF above 30), the median APC/IF is $277.67 with a range from $0 to $395.33. For the bin 2 journals (IF between 21 and 30), the median APC/IF rises to $427.59 with a range from $200.39 to $427.83. In contrast, for the bin 5 journals (IFs are less than 5), the median is $1051.11 with the widest range distribution ($0 to $2174.42), as seen in Figure 1C. In general, lower APC/IF values indicate higher impact-based cost effectiveness.

**Figure 1.**
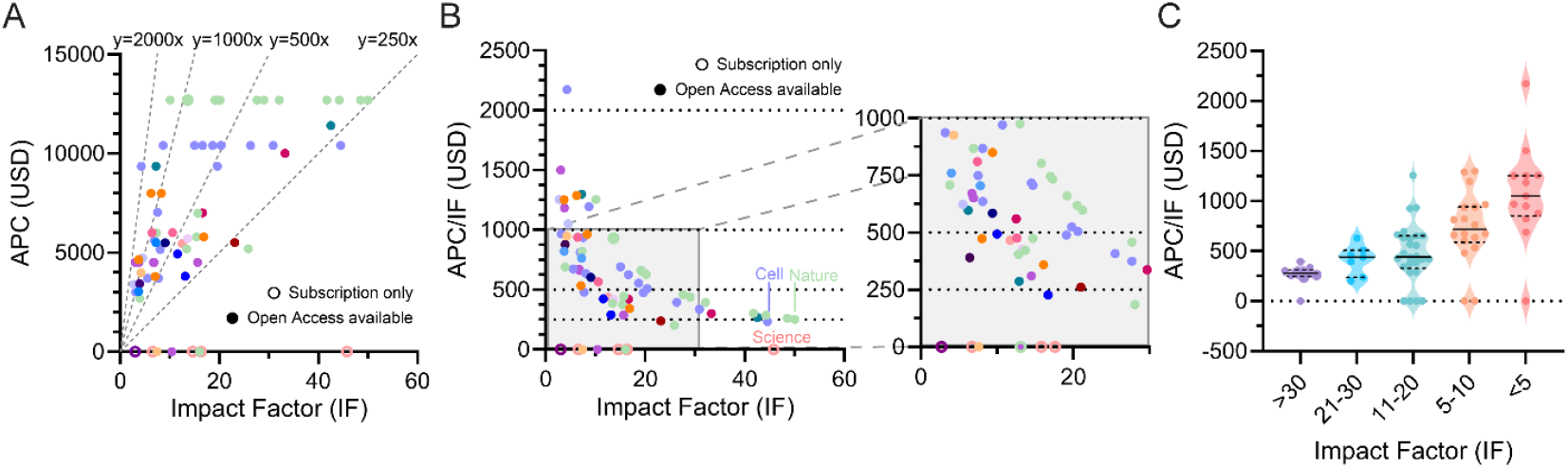
Cost-Impact evaluation of 74 representative biomedical journals using APC and IF. A. Article processing charge (APC) vs impact factor (IF) to illustrate the relationship and distribution between the publication cost and scientific impact of 74 representative biomedical journals. Each publisher of representative journals is represented by a different color. B. Impact-normalized processing charge (APC/IF) vs impact factor (IF) to show the impact-based cost effectiveness of different biomedical journals. C. Impact-normalized processing charge distribution based on five different impact factor bins.

While Journal Impact Factor is the most common journal impact metric, other metrics have been developed to address certain limitations of IF and provide additional insights. For example, CiteScore (CS) was developed by Scopus and is calculated by dividing the number of citations to documents published in a 4-year period by the number of documents in same 4-year period. The longer and symmetrical time-period of the calculation allows the sustained impact of a journal to be illustrated. Again, to understand if a journal’s scientific impact justifies its publishing cost, we performed a similar analysis of the relationship between a journal’s CiteScore and APC (Figure 2A and Supplemental Table 2). We also calculated the normalization of APC to the CS impact (APC/CS) vs CS allows authors to better evaluate the impact-based cost-effectiveness of different journals, as seen in Figure 2B. The APC/CS values showed similar patterns to APC/IF. The APC/CS ranges from 0 to 1395.52, with a median value of $346.05, and $218.90 of 25% percentile. The bin-based APC/CS analysis also revealed a reverse relationship between median APC/CS and CS values (Figure 2C). In bin 1 (CSs above 50), they have an APC/CS median of $171.36 and a range from $152.41 to $219.93. Bin 2 (CSs from 31 to 50) have an APC/CS median of $258.98 and range from $0 to $358.47. When the CS is less than 10 (bin 5), the APC/CS has a median of $715.12 and ranges from $0 to $1395.52. In general, we found APC/IF and APC/CS metrics are comparable. We found that 13 out 20 journals are ranked in the top 20 of both APC/IF and APC/CS.

**Figure 2.**
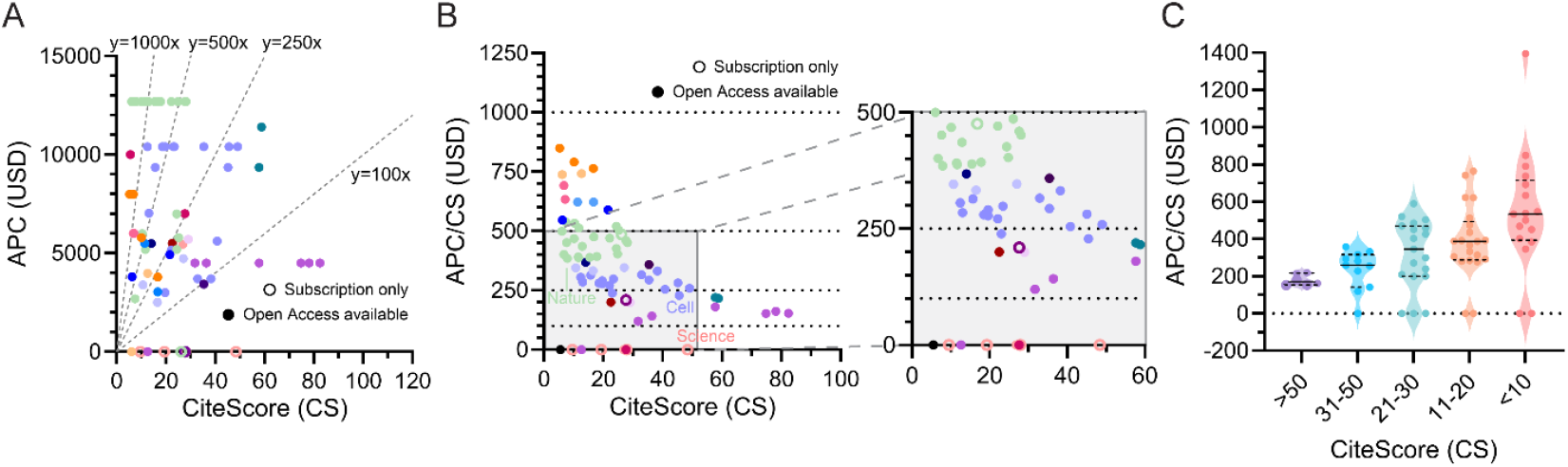
Cost-Impact evaluation of 74 popular biomedical journals using APC and CS. A. Article processing charge (APC) vs CiteScore (CS) to illustrate the relationship between a biomedical journal’s cost and scientific impact. Each publisher of our representative journals is represented by a different color. B. Impact-normalized processing charge (APC/CS) vs CiteScore (CS) to show the impact-based cost effectiveness of different biomedical journals. C. Impact-normalized processing charge distribution based on five different CS bins.

As APC/IF and APC/CS indexes are directly related to IF and CS, respectively, the impact-normalized indexes have the same intrinsic limitations as IF and CS (23). For example, IF and CS, as well as APC/IF and APC/CS, do not provide information about the quality or impact of individual articles published within that journal. Furthermore, because the citation practices and publication rates also differ significantly across disciplines, APC/IF and APC/CS cannot be used to compare journals from different fields. To mitigate this potential issue, we considered using Source Normalized Impact per Paper (SNIP) as an alternative metric for journal impact. SNIP enables direct comparison of journals in different subject fields. We computed APC/SNIP (Supplemental Table 3) and observed a similar pattern as other two CIE metrics (APC/IF and APC/CS). Finally, all CIE metrics (APC/IF, APC/CS, and APC/SNIP) have intrinsic biases toward journals that have high impact factors/cite scores. Therefore, as a guideline, we recommend using APC/CS or APC/IF cautiously for compatible journals in the same field (instead of different fields). Since all CIE metrics don’t have information of scope, researchers should use their own scientific judgement for selecting best fit journals or consult with journal editors for further information.

### Looking Forward

Scientific research stands as one of the most powerful drivers of advancements in medicine, technology, and public health. The shift toward open access publishing, while grounded in important ideals of transparency and public accessibility, has also introduced new financial challenges for researchers, particularly for labs and institutions with limited resources.

Sustaining open access for scientific publishing will require coordinated action across multiple stakeholders including government, funding agencies, publishers, and universities/institutions to work together to develop better and economic-efficient and sustainable models for scientific publishing. Solutions could include greater support for APCs from funding agencies and institutions, broader contribution to preprint and AAM repositories, expansion of “Green Open Access” options with a zero-embargo period from publishers.

In the interim, researchers need practical tools to navigate this complex publishing landscape. The Cost-Impact Effectiveness (APC/IF, APC/CS, APC/SNIP) metrics presented here are intended as one such resource—designed to help evaluate journals not only by scientific impact, but by the cost-effectiveness of their scientific impact. These data-driven metrics aim to support informed decision-making, particularly in environments where funding is limited and strategic publishing choices are essential. Next, we aim to create a tool to provide researchers with a convenient way to quickly look up the Cost-Impact Effectiveness metrics for journals of interest and evaluate the respective CIE metrics to help in the journal selection process. Looking ahead, the goal is not only to make science more open, but also to make publishing more cost-effective.

## Data Availability

Data is available on request.

## Acknowledgements

We would like to acknowledge the Scholarly Communications Librarians at the University of California San Diego Library, especially Karen Heskett, Allegra Swift and Teri Vogel of the UC San Diego Open Access Policy Team, for lending their expertise and providing the critical advice necessary to this commentary. This work was supported the Molecular Biophysics Training Grant (NIH T32 GM139795 to C.K.V.) and the Cancer Cell Signaling and Communication Training Program (NIH T32CA009523 to J.C.H.).

## Author Contributions

J.C.H. compiled the data, analyzed the data and made the figures. D.W. oversaw the data collection and analysis. J.C.H. and C.K.V. wrote the manuscript. J.C.H., C.K.V., and D.W. edited the manuscript.

---

**Supplemental Table 1.**
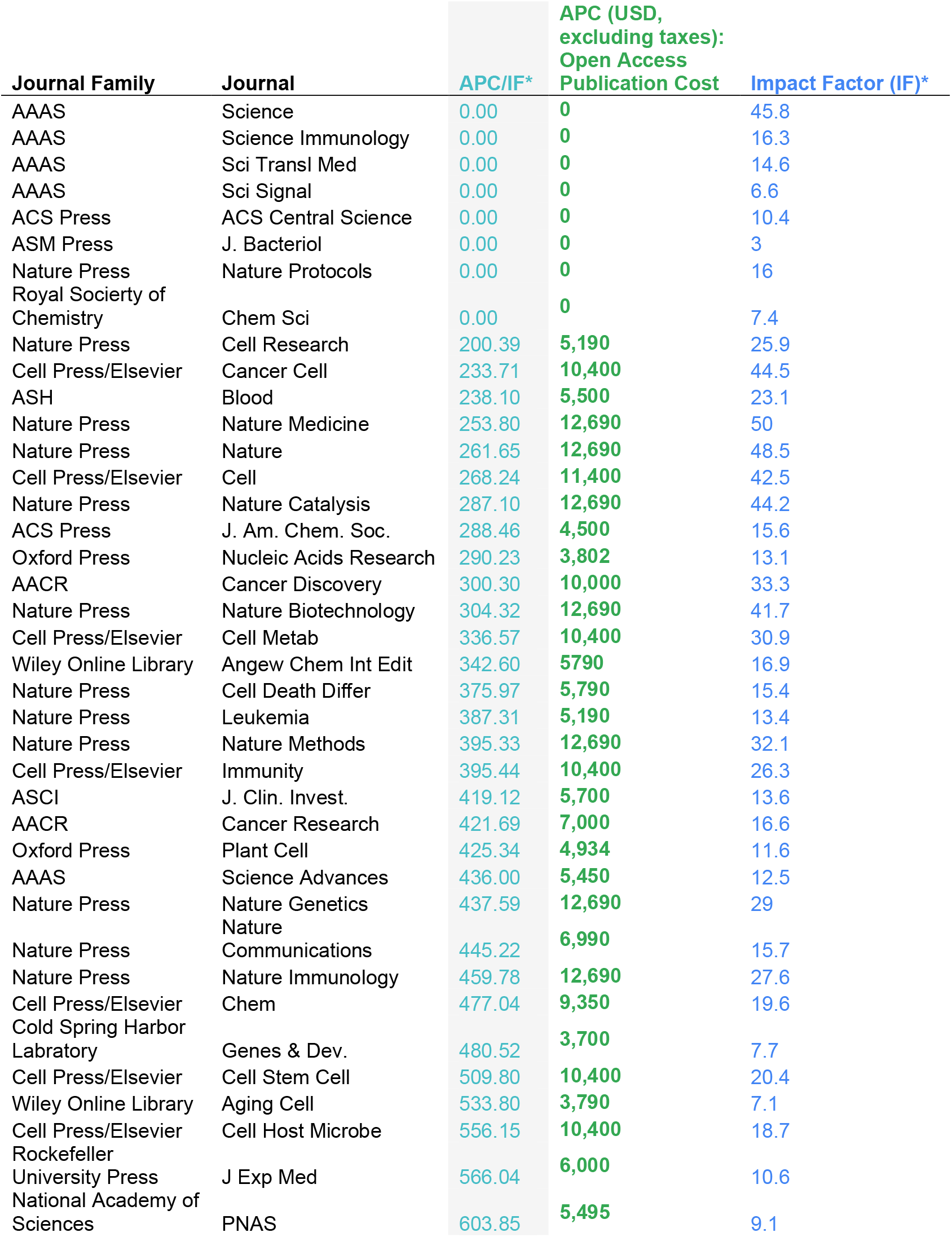
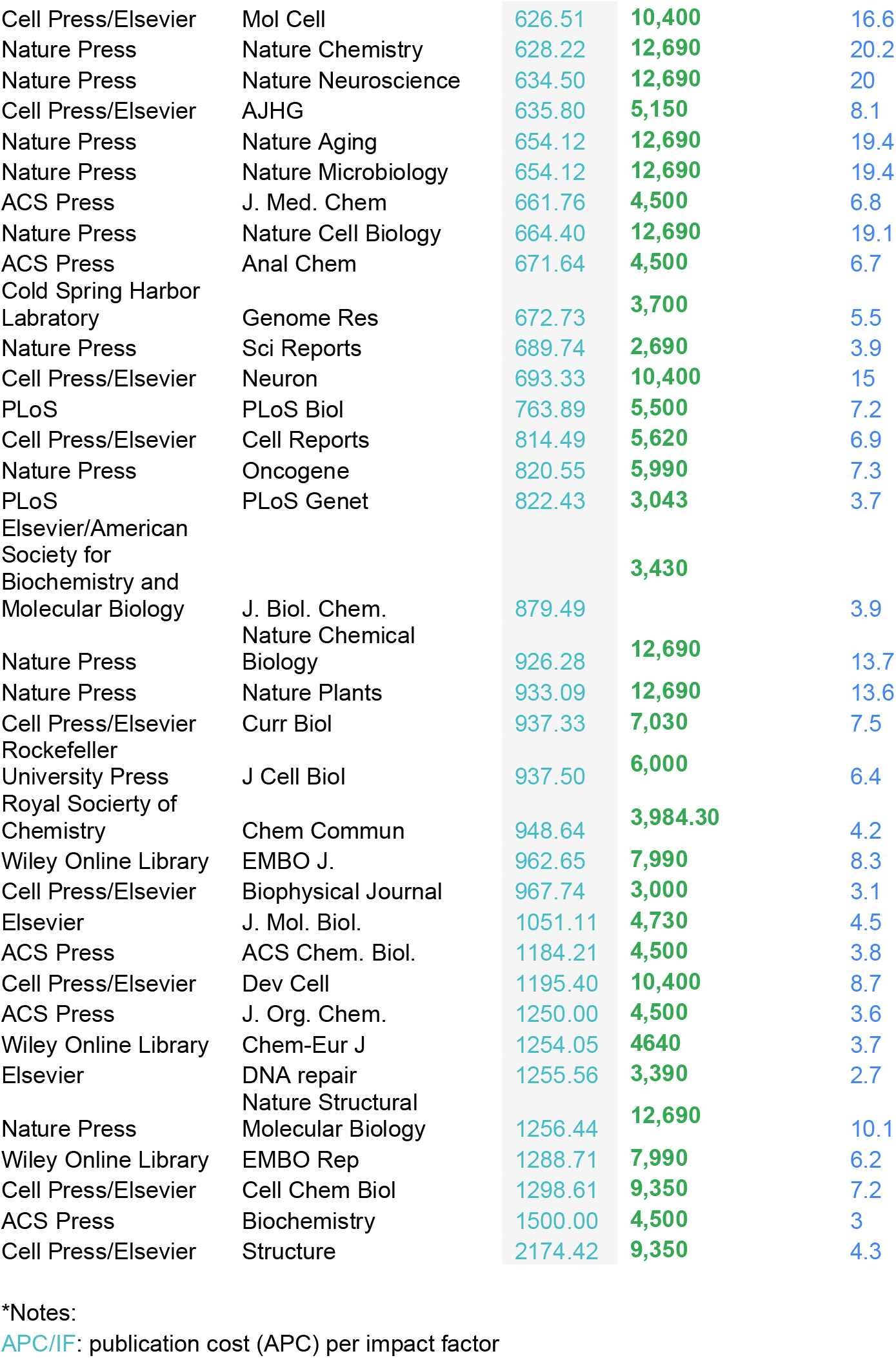
Cost-Impact Effectiveness Metric APC/IF. The journal publication cost (APC) normalized to impact factor (APC/IF) per journal, sorted from lowest APC/IF to highest APC/IF.

**Supplemental Table 2.**
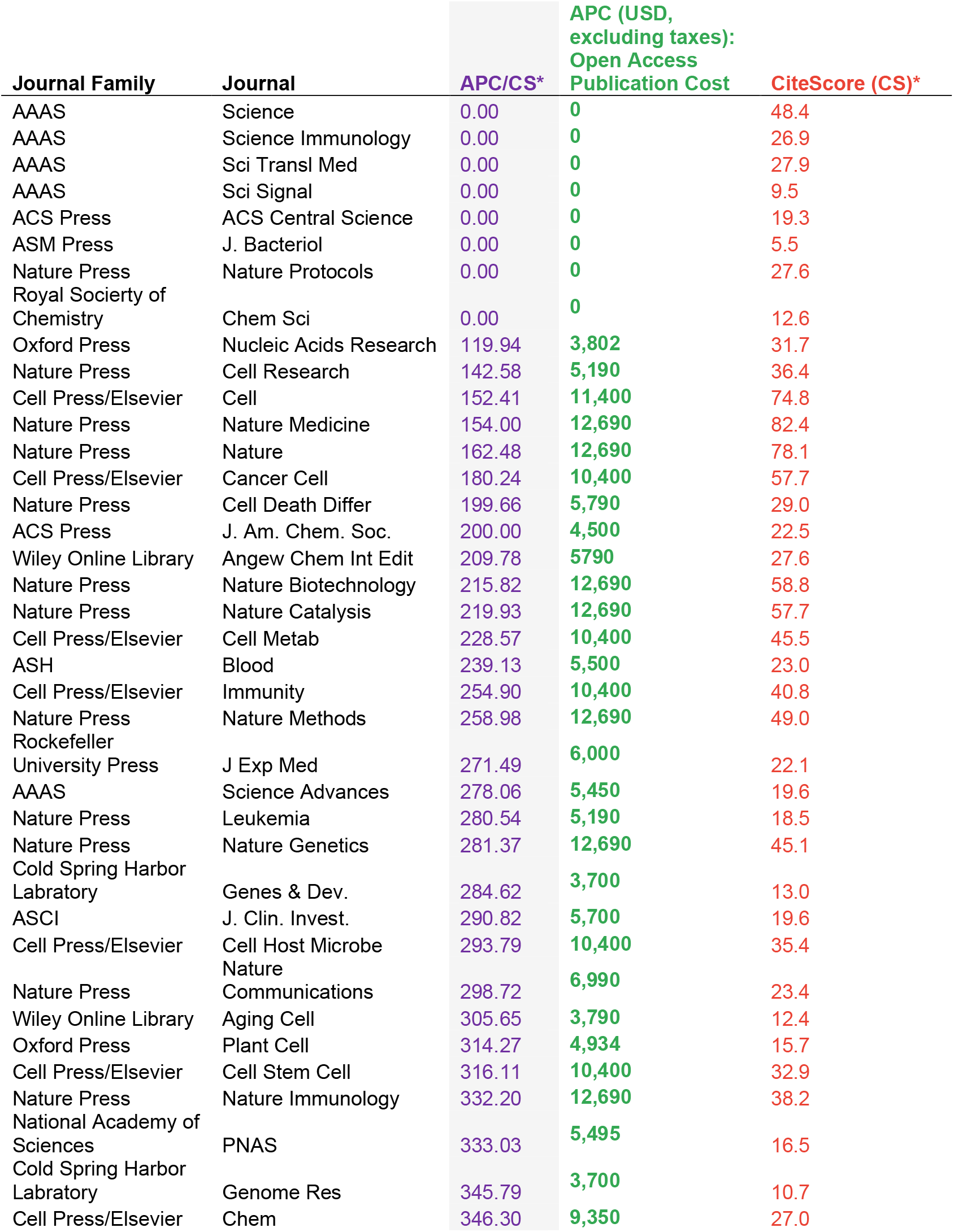
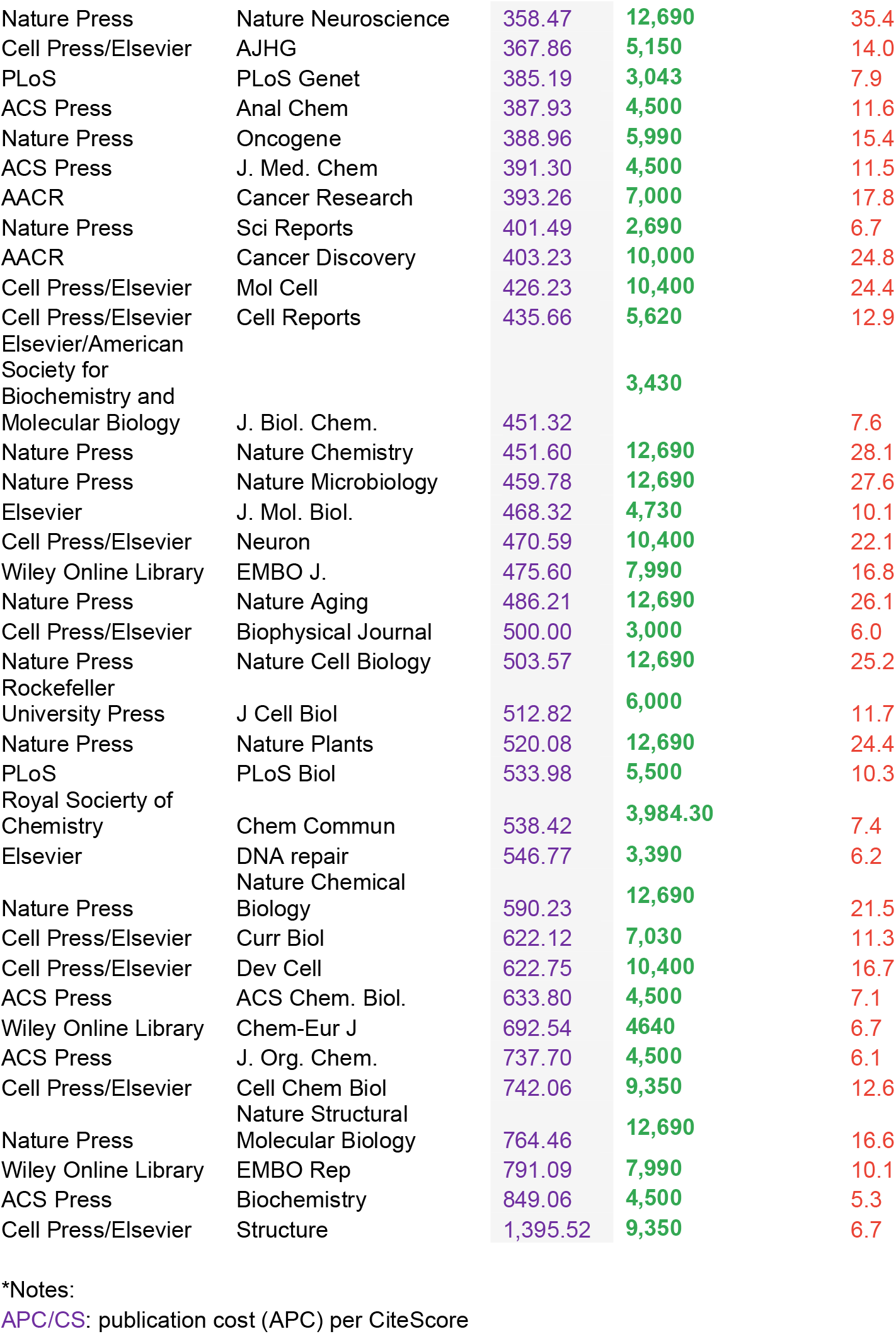
Cost-Impact Effectiveness Metric APC/CS. The journal publication cost (APC) normalized to CiteScore (APC/CS) per journal, sorted from lowest APC/CS to highest APC/CS.

**Supplemental Table 3.**
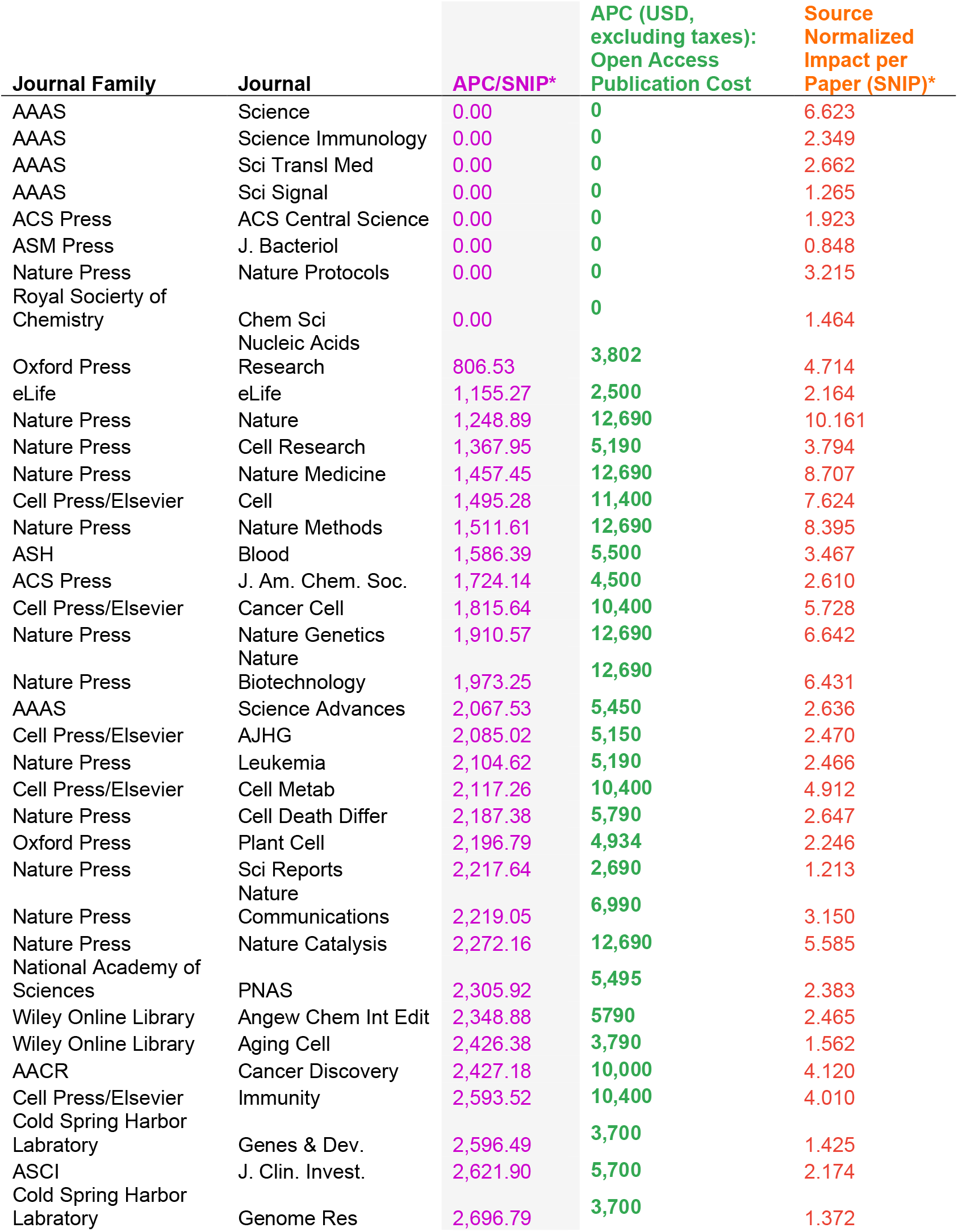
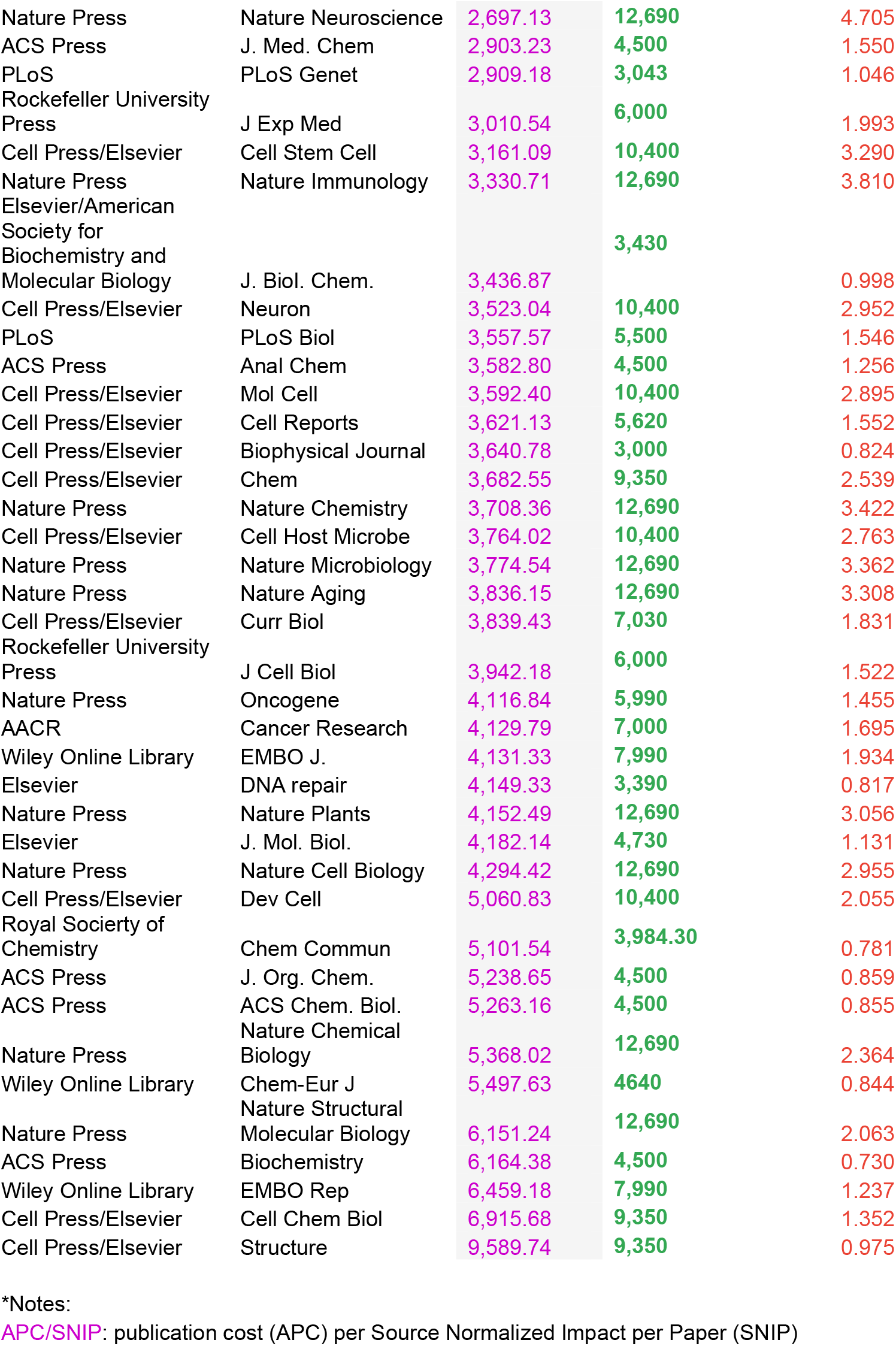
Cost-Impact Effectiveness Metric APC/SNIP. The journal publication cost (APC) normalized to Source Normalized Impact per Paper (APC/SNIP) per journal, sorted from lowest APC/SNIP to highest APC/SNIP.

